# Single molecule analysis of lamin dynamics

**DOI:** 10.1101/371104

**Authors:** Leonid A. Serebryannyy, David A. Ball, Tatiana S. Karpova, Tom Misteli

## Abstract

The nuclear envelope (NE) is an essential cellular structure that contributes to nuclear stability, organization, and function. Mutations in NE-associated proteins result in a myriad of pathologies with widely diverse clinical manifestations, ages of onsets, and affected tissues. Notably, several hundred disease-causing mutations have been mapped to the *LMNA* gene, which encodes the intermediate filament proteins lamin A and C, two of the major architectural components of the nuclear envelope. However, how NE dysfunction leads to the highly variable pathologies observed in patient cells and tissues remains poorly understood. One model suggests alterations in the dynamic properties of the nuclear lamina and its associated proteins contribute to disease phenotype. Here, we describe the application of single molecule tracking (SMT) methodology to characterize the behavior of nuclear envelope transmembrane proteins (NETs) and nuclear lamins in their native cellular environment at the single molecule level. As proof-of-concept, we demonstrate by SMT that Halo-tagged lamin B1, Samp1, lamin A, and lamin A Δ50 have distinct binding and kinetic properties, and we identify several disease-relevant mutants which exhibit altered binding dynamics. SMT is also able to separately probe the dynamics of the peripheral and the nucleoplasmic populations of lamin A mutants. We suggest that SMT is a robust and sensitive method to investigate how pathogenic mutations or cellular processes affect protein dynamics at the NE.

## 1. Introduction

The nucleus is enclosed by a bi-layered nuclear envelope (NE) and an underlying proteinaceous lamina network [1, 2]. The outer nuclear membrane (ONM) is contiguous with the endoplasmic reticulum (ER) and separated from the inner nuclear membrane (INM) by the perinuclear space. These two membranes are connected by nuclear pore complexes (NPCs). Whereas NPCs facilitate protein transport between the nucleus and cytoplasm, proteins exchange between the ONM and INM via diffusion within the lipid bilayer [3, 4]. Despite the large number of proteins found at the NE, the two nuclear membrane leaflets maintain a preferential protein composition of NE transmembrane proteins (NETs) [4-7]. The preferential localization of NETs to the ONM or INM is likely due to compartment specific binding partners [4]. Below the INM lies the nuclear lamina, a 3.5 nm network of tetramers of type V intermediate proteins that maintain nuclear structure, function, and the composition of the NE [8-12]. The nuclear lamina is composed of A- and B-type lamins that are differentially expressed among cell types as well as a host of lamin-associating proteins. In humans, the *LMNA* gene encodes the two major A-type lamins, lamin A and C, while the *LMNB1* and *LMNB2* genes encode the two major B-type lamins, lamin B1 and B2. Together with the nuclear lamina, NETs have been found to be major regulators of chromatin organization, mechano- and signal transduction [12, 13].

The functional importance of the NE is impressively illustrated by the fact that mutations in NE-associated proteins result in over 20 distinct human pathologies [5, 9, 14, 15]. The majority of these mutations occur in the *LMNA* gene. Notably, >300 disease-related mutations have been mapped to *LMNA* and lead to a highly diverse and tissue-specific set of pathologies including striated muscle dysfunction [dilated cardiomyopathy (DCM), congenital muscular dystrophy (CMD), limb-girdle muscular dystrophy (LGMD1B), Emery-Dreifuss muscular dystrophy (EDMD)], adipose and skeletal tissue defects [Dunnigan-type familial partial lipodystrophy (FPLD) and Mandibuloacral dysplasia (MAD)], an axonal myelination neuropathy [Charcot-Marie-Tooth syndrome type 2b (CMT2B1)], and variable multisystem progeria syndromes characterized by accelerated aging with growth retardation, metabolic dysfunction, and vascular abnormalities [Werner syndrome (WRN), Hutchinson-Gilford progeria syndrome (HGPS)] [2, 9, 10, 14]. Mutations across the *LMNA* gene may manifest as similar pathologies, and cases have been identified where different mutations at the same residue can lead to strikingly different diseases. For example, mutation of residue 527 from an arginine to a proline has been linked to EDMD, while conversion to a histidine has been implicated in MAD, and a mutation to a cysteine correlates with progeria and MAD phenotypes. Similarly, mutation of the neighboring residue 528 from a threonine to a lysine or arginine is linked to EDMD, whereas a change to a methionine results in FPLD [9].

To explain the pathological diversity arising from NE dysfunction, a number of non-mutually exclusive hypotheses have been suggested; notably, that these diseases are a result of mechanical dysfunction and uncoupling between the cytoskeleton and nucleus via the linker of nucleoskeleton and cytoskeleton (LINC) complex, a consequence of altered nuclear signaling and gene expression, or the premature depletion of regenerative stem cells [2, 9]. However, these mechanisms have been unable to fully explain the complex pathologies that arise from NE dysfunction, pointing to the need for new approaches to characterize the molecular properties of lamins and NETs.

Here, we apply single molecule tracking (SMT) to quantitatively assess lamin and NET dynamics. A hallmark of SMT is its ability to characterize the trajectories of individual molecules with nanometer spatial resolution and millisecond temporal precision in the context of living cells [16, 17]. Like other live-cell imaging methods, such as fluorescence correlation spectroscopy (FCS) and fluorescence recovery after photobleaching (FRAP), SMT employs an exogenously expressed protein fused to a fluorophore to measure protein dynamics. However, SMT offers the advantages of higher spatial resolution, discrimination between heterogeneous populations of molecules in the same cell, and is able to more easily assess the dynamics of relatively immobile molecules [17]. Previous studies have utilized SMT to study the dynamics of mRNAs, chromatin remodelers, and transcription factor binding events, among others [16-23]. We use highly inclined and laminated optical sheet (HILO) illumination, sub-optimal transient transfection of Halotag-fused lamins and NETs, and low labelling concentrations of a Halotag ligand to perform time-lapse imaging of a variety of NE-associated proteins to determine the fractions of bound and unbound molecules as well as dwell times. We show that SMT is sufficiently sensitive to discriminate between dynamics of different lamins, lamin mutants and NETs and the observed kinetics are in line with known biology. In addition, we apply SMT to a library of lamin A mutants to characterize the link between pathological mutations in lamin A with their respective dynamic behavior.

## 2. Methods

### 2.1. Cell lines and cell culture

U2OS cells were cultured in DMEM (Thermofisher Scientific) supplemented with 10% FBS (Sigma-Aldrich), L-glutamine, and penicillin/streptomycin (Thermofisher Scientific). To create lamin B receptor (LBR) fragment-EGFP stably expressing cells, U2OS cells were transiently transfected with pcDNA-LBR-EGFP-polyA (Addgene plasmid # 59748) for 48 h using Lipofectamine 2000 (Thermofisher Scientific) according to the manufacturer’s instructions before addition of 1 mg/mL G418 (Invivogen) to the media. Cells were then cultured in the presence of G418 for several weeks until only EGFP positive cells remained in the population. This line of U2OS cells was subsequently used for transient transfections of Halo-tagged proteins.

To prepare cells for SMT, U2OS cells were seeded into two-well Lab-Tek chamber slides (Thermofisher Scientific) for at least 16 h. Cells were then transiently transfected overnight using 2 μL Lipofectamine 2000 and 0.5 μg of DNA per chamber in OptiMem (Thermofisher Scientific). After 16–48 h, cells were incubated with 1 nM JF646 for 30 min at 37 °C [24]. Cells were subsequently washed in PBS for 5 min for three cycles. The remaining PBS was aspirated, and cells were incubated in Fluorobite DMEM (Thermofisher Scientific) supplemented with 10% FBS, L-glutamine, and penicillin/streptomycin. Where indicated, 1 μM lonafarnib, a farnesyltransferase inhibitor (FTI) was added after transfection.

### 2.2. Plasmids

The N-terminal Halotag fragment was derived from pHalotag-CREB, a kind gift from Dr. Gordon Hager (LRBGE, NIH, Bethesda, MD), and purified using the *NheI* and *XhoI* restriction enzymes. Halotag was used in exchange for the N-terminal fluorophore in pEGFP-C1, pEGFP-lamin A, pEGFP-lamin A Δ50 [25], mWasabi-LaminB1-10 (Addgene plasmid # 56507), and YFP-Samp1 (Spindle Associated Membrane Protein 1) which was a gift from Dr. Einar Hallberg (Stockholm University, Stockholm, Sweden).

Halo-lamin A mutations were created using the QuikChange XL Site-Directed Mutagenesis Kit according to the manufacturer’s instructions (Agilent). The primers used for mutagenesis and the resulting pathologies from the individual mutations are listed in **Supplementary Table 1**.

### 2.3. Immunofluorescence staining

To prepare the cells for confocal imaging, ~60,000 U2OS cells were seeded on 12 mm glass coverslips (Thermofisher Scientific) in 24-well plates (Denville Scientific). To prepare cells for high-throughput imaging, 5,000 cells were seeded into clear bottom 96-well plates (Perkin-Elmer). Cells were transiently transfected with Halo-lamin A or the lamin A mutants 24 h after plating. After 24–48 h, cells were rinsed in PBS, fixed in 4% paraformaldehyde (Electron Microscopy Sciences) for 10 min, washed in PBS, and then permeabilized with 0.3% Triton X100 (Sigma-Aldrich) for 10 min. Cells were subsequently washed in PBS and blocked in 2% BSA (Sigma-Aldrich) for 1 h at room temperature. To label cells, they were incubated with a polyclonal anti-Halotag antibody (Promega) at a 1:200 dilution in 2% BSA for 1 h at room temperature. Cells were then washed 3 times for 10 min in PBS and incubated with Alexa Fluor 594 labeled donkey anti-rabbit secondary antibody (Thermofisher Scientific) at 1:400 in 2% BSA for 1 h at room temperature in the presence of 2 μg/mL DAPI. Cells underwent another round of washes in PBS before mounting on coverslips with Vectashield (Vector Labs) or high-throughput imaging. Similar staining was observed when cells were directly labeled with 3 μM JF646.

### 2.4. HILO microscopy

Single molecule tracking (SMT) was performed at the NIH/NCI CCR/LRBGE Optical Microscopy Core (Bethesda, MD, USA). A custom inverted wide-field microscope equipped with a 100x oil immersion objective (NA = 1.49, Olympus), and three EMCCDs (Andor iXon 888) for simultaneously three color imaging based on designs previously described [26, 27] was used for HILO excitation of the sample [17] with solid-state lasers at wavelengths of 488, 561, and 647 nm (Coherent OBIS). Fluorescence emission is separated from scattered laser light by use of a quad-band dichroic (ZT405/488/561/647, Chroma Technology Corp., VT, USA), and the emission bands are separated by two long-pass filters (T588lpxr, T660lpxr) and emission filters (525/50, 609/58 and 736/128, Semrock, Inc., NY, USA). A temperature- and CO_2_-regulated stage was used to maintain the cells at 37 °C and 5% CO_2_. Images were acquired with 30 ms exposure times at 200 ms intervals for a total of approximately three minutes using a 488 nm (100 μW) excitation source to image LBR fragment-EGFP as a marker of the nuclear periphery and a 647 nm (2 mW) excitation source to image JF646-labeled Halotag.

### 2.5. SMT analysis

Molecule tracking was performed using ‘TrackRecord’ software developed in Matlab as previously described [17]. A maximum projection of the LBR Fragment-EGFP image stack was used to create a region of interest either selecting the entire nucleus or segmenting the nuclear periphery from the nuclear interior. Images were filtered for potential molecules by applying Wiener, top-hat, and size filters to reduce speckle noise, correct for uneven illumination, and highlight features of approximately 5 pixels in area. Only the brightest pixel is kept when multiple molecules are found within 7 pixels of each other. Each seed is fitted to a two-dimensional Gaussian to determine molecule position, and trajectories are created using a nearest-neighbor algorithm. A maximum movement of 4 pixels per frame was set and only tracks at least 6 frames long were analyzed. To account for fluorophore blinking, single-frame gaps in trajectories were filled with the average position of the molecule in the flanking frames. Typically, approximately 500,000 molecules were analyzed per Halo-tagged protein. Segments of tracks were considered bound only if their frame-to-frame displacement did not exceed a distance of 220 nm for four frames, which reduces the probability to 0.002 that a diffusing molecule will be considered bound [28]. The total bound fraction was then calculated as the ratio of the number of bound track points to the total number of molecules. The duration of the bound track segments was used to generate a survival distribution, which is the complement to the cumulative distribution function (1-CDF). The survival distribution was then corrected for photobleaching by dividing it by the decay in the number of molecules found over time, normalized to the bound fraction and fit to a bi-exponential curve by least square to extract the mean residence times, and proportions of fast- and slow-bound molecules. Statistical analysis was performed between confidence intervals among each condition using Z-tests.

### 2.6. Confocal microscopy

Confocal microscopy was performed using a Zeiss LSM 780 laser scanning microscope equipped with a 63x objective (NA = 1.4) and a 561 nm excitation source. Images were acquired at the NIH/NCI CCR/LRBGE Optical Microscopy Core (Bethesda, MD, USA).

### 2.7. High-throughput imaging

High-throughput confocal imaging was performed using a 63x water objective on a Yokagawa CV7000 spinning disk microscope at the CCR High-throughput Imaging Facility (NIH, Bethesda, MD, USA). For excitation, two laser lines were used (405 nm and 561 nm), and images were acquired with 0.5μm-1.0 μm Z-sections for at least 10 randomly chosen fields of view in an automated manner. >50 cells were analyzed over 3 experiments. Images were binned 2×2 and average fluorescence intensities from the Z-stacks were saved. Images were processed using Columbus software (Perkin-Elmer) to identify and segment nuclei using the DAPI nuclear stain. Nuclear morphology and mean fluorescence intensity per well was calculated using anti-Halotag staining.

## 3. Results

### 3.1. Single molecule tracking methodology

Tracking individual molecules within cells enables real-time interrogation of protein behavior in a cellular context [16], however, several requisites must be met. A bright and stable fluorophore used at a low labeling density is required to discriminate and track individual molecules over an extended period. In conjunction, a highly sensitive camera that is able to detect individual molecules as well as a microscope setup to optimize the signal-to-noise ratio and limit out-of-focus fluorescence are critical. Therefore to study lamin and NET dynamics, we used a custom HILO imaging system with three electron-multiplying charge-coupled device (EMCCD) cameras similar to the configuration previously used to perform SMT on transcription factors [17, 18]. HILO microscopy employs an inclined incident laser beam at the periphery of the objective lens and generates a thin optical light sheet that reduces background compared to epi-fluorescent illumination and therefore increases the signal to background ratio of the image [29].

To accurately mark the nuclear periphery as a region of interest (ROI) for downstream image analysis in subsequent SMT experiments, we generated U2OS cells stably expressing an EGFP-tagged N-terminal fragment of the Lamin B Receptor (LBR), which has previously been shown to localize to the nuclear periphery [30, 31]. Alternatively, we found that the nuclear periphery could also be accurately defined in our experiments by using maximum intensity projections of the Halo-tagged proteins analyzed. U2OS cells stable expressing LBR fragment-EGFP were seeded in glass bottom chambered slides, allowed to adhere overnight, and transiently transfected with Halo-tagged proteins (**Figure 1**). Before imaging, U2OS cells were incubated with the Halo ligand JF646 at 1 nM for 30 min at 37 °C, washed 3 times 5 m in PBS, and placed in phenol-free media (see Methods). The heterogeneous expression of Halotag after transient transfection facilitated easier selection of cells with a labeling density suitable for the identification of single molecules. Cells were imaged with 30 ms exposure times at 200 ms intervals for 900 acquisitions using simultaneous 488 nm and 647 nm excitation. We found images acquired every 30, 200, and 1000 ms provided similar results in Halo-lamin A expressing cells (data not shown). The 200 ms imaging interval was chosen to balance photobleaching but still capture the population of molecules that exhibit fast dynamics. Images were acquired from at least 10 cells among two independent experiments and typically ~500,000 molecules were tracked for quantitative analysis per protein; we found little variation between experiments and sample size.

**Figure 1:**
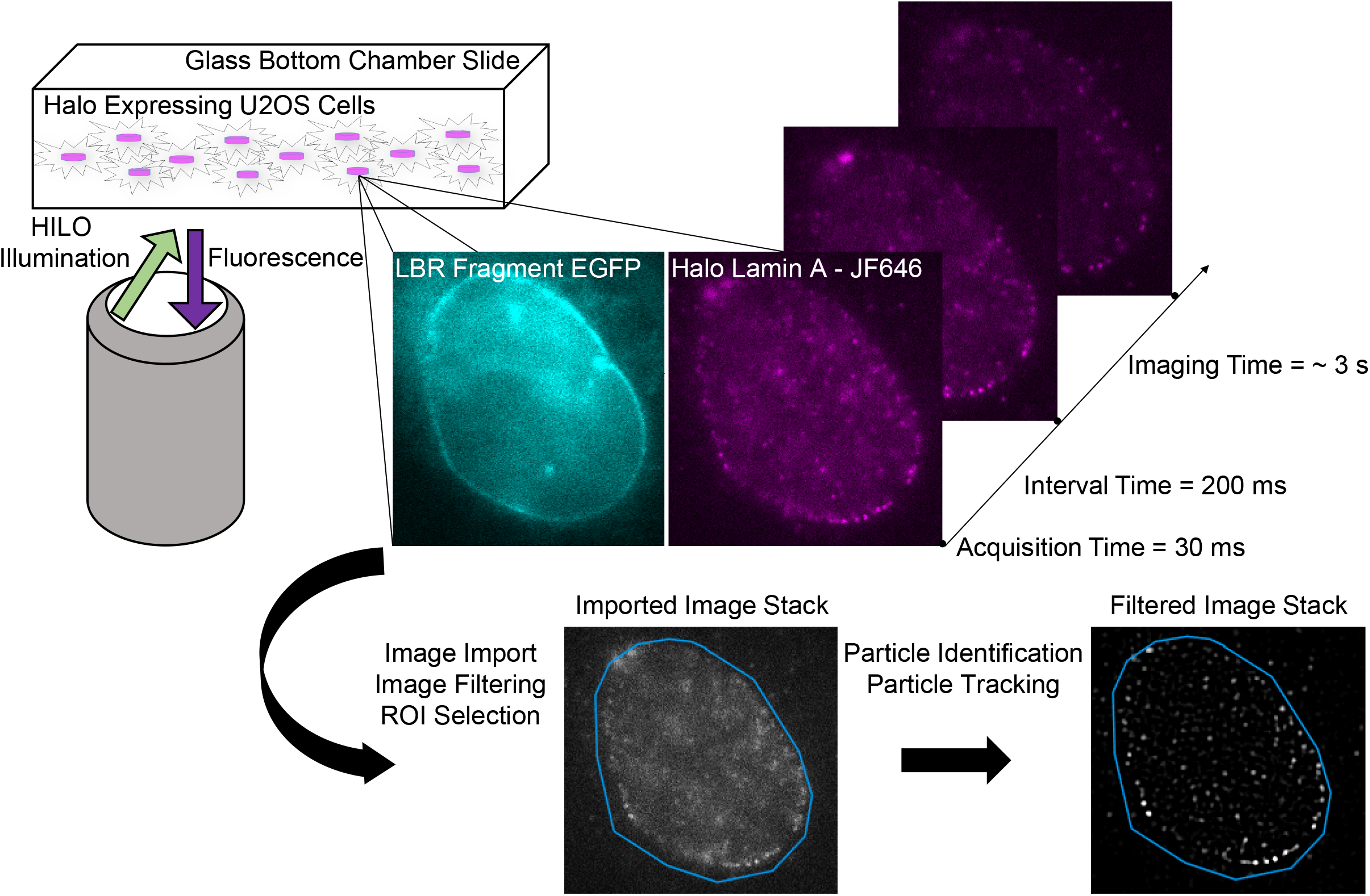
Single-molecule tracking (SMT) methodology. U2OS cells stably expressing an EGFP-tagged N-terminal fragment of Lamin B Receptor (LBR) to mark the nuclear periphery are grown on glass bottom chamber slides. Once at an optimal confluency, cells are transiently transfected with a Halo-tagged protein of interest. After 24 h of transfection, 1 nM of Halotag ligand (JF646) is added for 30 min followed by a series of washes to remove unbound ligand before imaging. Highly inclined and laminated optical light sheet illumination (HILO) is employed to image a thin optical section and limit out-of-focus fluorescence. A series of images are taken in the same plane using simultaneous 488 nm and 647 nm excitation with a 30 ms acquisition time every 200 ms. Following acquisition, imaging channels are split using Image J and imported into a customized MATLAB software to annotate a region of interest (i.e. LBR fragment localization), individual spots are then tracked over time. Single-molecule tracks collected amongst different cells and experiments are aggregated, undergo photobleaching correction, and are fit to a one-, bi-, or multi-component exponential distribution.

After image acquisition, image stacks were imported into the ‘TrackRecord’ Matlab software as previously described [17]. Images were filtered to remove speckle noise, to correct for uneven HILO illumination, and to enrich for trackable molecules. ROIs were then chosen based on LBR fragment-EGFP fluorescence and molecules were fit to a Gaussian distribution. Trajectories were calculated using a nearest neighbor algorithm with parameters set to include molecules that move a maximum of four pixels in one frame and are tracked for at least six frames, allowing for a gap of one frame in the trajectory. The survival distribution was obtained from bound track segments (as defined in Methods) in each condition and was fit to a bi-exponential decay after correcting for photobleaching and normalizing to the total bound fraction. Therefore, the analysis results in three distinct populations with unique residence times: diffusing, fast-bound, and slow-bound molecules (**Figure 2**).

**Figure 2:**
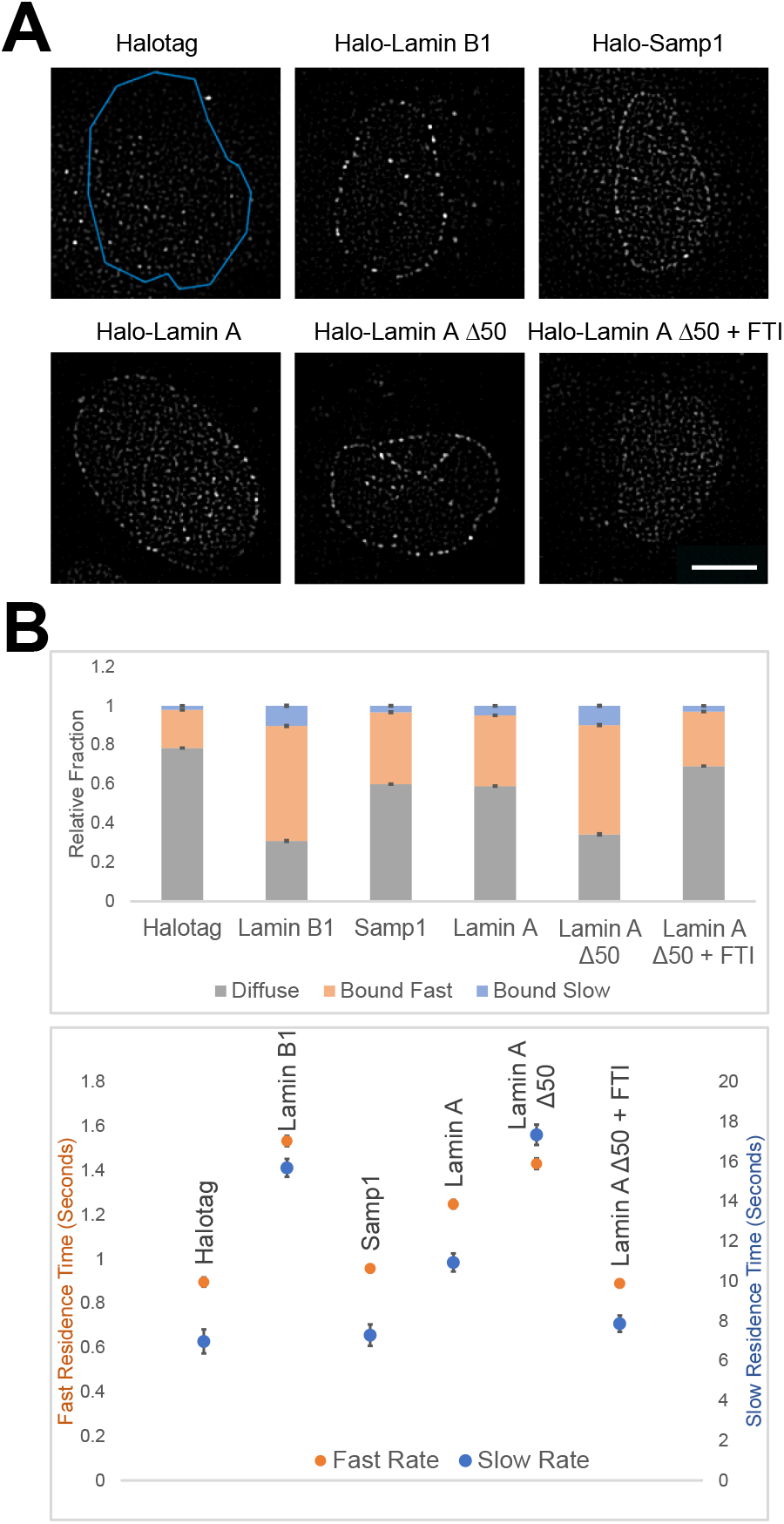
SMT of lamins and NETs at the nuclear periphery. (A) Representative images after post-processing of U2OS cells transfected with Halotag alone, Halo-lamin B1, Samp1, lamin A, lamin A Δ50, and lamin A Δ50 treated with 1 μM farnesyltransferase inhibitor for 16 h. (B) SMT of the cells per condition was compiled and nuclear molecule tracks were fit to a bi-exponential distribution. Quantification of molecule tracks indicates the ratio of identified molecules that are diffuse (grey), fast bound (orange), and slow bound (blue; top) as well as the associated dwell times (bottom). Halo-tagged proteins show increased fractions of bound molecules and dwell times as compared to non-specific binding of Halotag alone. Exposure time 30 ms; acquisition time 200 ms; 900 images per cell. Values represent means and 95% confidence intervals of at least 10 cells and 2 experiments. Scale bar = 10 μm.

### 3.2. Single molecule tracking of proteins at the nuclear periphery

As proof of principle, we assayed the dynamics of a variety of lamins and NETs that exhibit different levels of association with the nuclear periphery (**Figure 2A**). As a negative control SMT of Halotag alone resulted in a highly diffuse population (78%) with short dwell times for the remaining fraction of fast bound molecules (20%, 0.90 sec), and a very minor slow bound fraction (2%, 7.0 sec), which may represent diffusing Halotag molecules transiently trapped in higher order chromatin features such as heterochromatin (**Figure 2B**). In contrast, analysis of lamin B1, which is known to form stable polymers and is physically incorporated into the INM [9, 32], showed that 59% of molecules had a fast bound dwell time of 1.5 sec and 10% of molecules had a slow bound dwell time of 15.7 sec, considerably longer than those of Halotag alone (p < 0.0001 for all conditions). Samp1, an INM transmembrane protein and component of the LINC complex [33, 34], was predominantly localized to the nuclear periphery with some nucleoplasmic and ER fluorescence (**Figure 2A**). Samp1 exhibited faster dwell times and a smaller bound fractions than lamin B1 (Samp1: 40% vs. lamin B1: 69%; p < 0.0001 across all parameters). Interestingly, Samp1 exhibited a similar, yet statistically different, proportion of bound molecules as its binding partner lamin A (44% bound; p < 0.01), albeit with faster dwell times (Samp1: fast bound dwell time of 0.96 sec, slow bound dwell time of 7.3 sec vs. lamin A: fast bound dwell time of 1.2 sec, slow bound dwell time of 12 sec; p < 0.0001) (**Figure 2B**). The proportion of bound lamin A molecules and the associated dwell times are reflective of the distribution of lamin A between a well-established dynamic nucleoplasmic population as well as a stably polymerized population at the nuclear periphery [32, 35, 36] (**Figure 2B**). We conclude that SMT is able to sensitively distinguish binding dynamics of NE-associated proteins.

To assess if SMT is able to detect changes in dynamics caused by individual protein modifications, we expressed a lamin A construct with an internal deletion of 50 amino acids near its C-terminus (Halo-lamin A Δ50). Lamin A Δ50, also referred to as progerin, is the causal driver of the pre-mature aging disorder Hutchinson-Gilford Progeria Syndrome (HGPS) [25, 37-40]. Due to the internal deletion, progerin remains permanently farnesylated and is stably integrated into the INM [9, 41, 42]. By SMT analysis, lamin A Δ50 exhibited larger bound fractions compared to wild type lamin A (fast bound: 55.9% vs. 35.2%, p < 0.0001; slow bound: 9.8% vs. 6.7%, p < 0.0001) and longer dwell times (fast bound dwell time: 1.4 sec vs. 1.2 sec, p < 0.0001; slow bound dwell time: 17.4 sec vs. 11.9 sec, p < 0.0001) (**Figure 2B**). Like lamin B1, lamin A Δ50 is able to polymerize and is physically bound to the INM [42]. These properties likely explain the more comparable behavior measured by SMT between lamin B1 and lamin A Δ50 with 58.8% vs. 55.9% of the tracked molecules to be fast bound (p = 0.174), 9.8% vs. 10.3% slow bound (p < 0.0001), fast bound dwell times of 1.5 sec vs. 1.4 sec (p < 0.0001), and slow bound dwell times of 16 sec vs. 17 sec (p < 0.0001), respectively (**Figure 2B**). To ensure our SMT measurements were reflective of biological function, we expressed lamin A Δ50 in the presence of the farnesyl transferase inhibitor (FTI) lonafarnib, which prevents lamin A Δ50 farnesylation and thus integration into the INM [41]. Farnesyltransferase inhibition dramatically reduced the percentage of bound molecules of lamin A Δ50 as compared to untreated lamin A Δ50 cells (fast bound fraction from 55.9% to 27.9%, slow bound fraction from 9.8% to 2.9%, fast bound dwell time from 1.43 sec to 0.89 sec, and the slow bound dwell time from 17.4 sec to 7.8 sec; p < 0.0001 for all conditions) (**Figure 2B**). Taken together, these data demonstrate the ability of SMT to extract information about localization as well as binding dynamics with the sensitivity to distinguish distinct modes of interactions of NE-associated proteins.

### 3.3. Mapping dynamic properties of lamin A mutants

We next used SMT to characterize the potential effects of point mutations in lamin A on the proteins’ dynamic properties. In addition, we sought to investigate if correlations exist between disease-specific mutations and their dynamics. Therefore, we generated a series of mutations in the Halo-lamin A construct that have been shown to cause a spectrum of laminopathies, particularly mutations associated with CMD, EDMD, MAD, FPLD, WRN, and progeria (**Figure 3A, Supplementary Table 1**). To exclude the possibility that the observed changes in dynamics are due to aberrant nuclear morphology induced by the mutants, we transiently expressed the lamin A mutants in U2OS cells. High-throughput imaging and subsequent image analysis revealed no obvious changes in nuclear area upon expression of any of the mutants (**Figure 3B**). Furthermore, scoring for dysmorphic nuclei revealed a clear phenotype in only Halo-lamin A Δ50 expressing cells as previously reported (**Figure 3C**) [37, 40].

**Figure 3:**
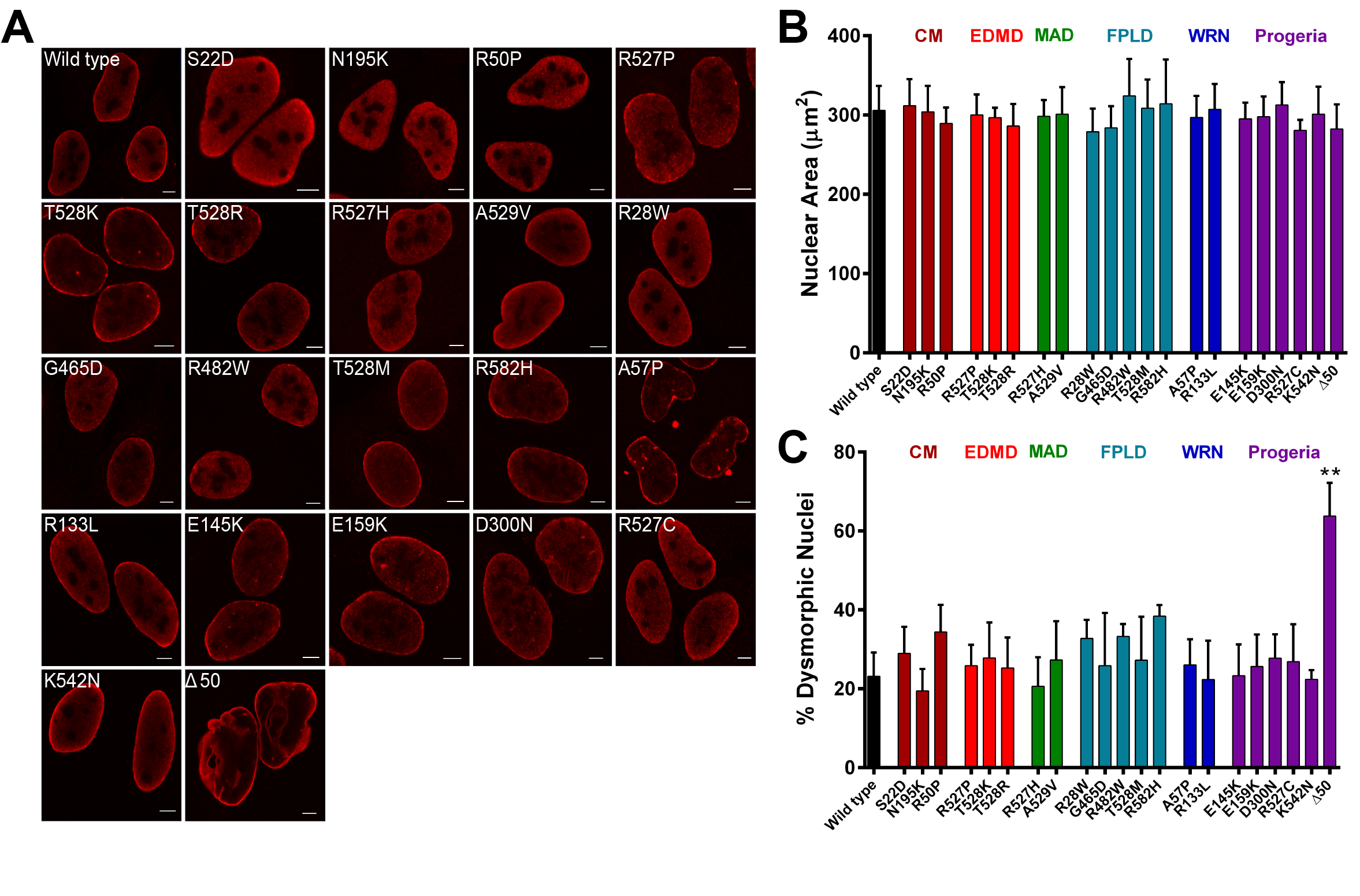
Morphology of nuclei expressing Halo-lamin A mutants. (A) Confocal imaging of U2OS cells expressing the indicated Halo-lamin A mutants and labeled with anti-Halo antibody (red). Scale bars = 5 μm. (B) Nuclear area (μm^2^) of U2OS cells transiently expressing the indicated Halo-lamin A mutants. Mutations are categorized by the characteristic disease outcome in human patients and by residue position. Mean ± SEM; N = 3 experiments and > 50 cells per condition. CM, cardiomyopathy defect; EDMD, Emery-Dreifuss muscular dystrophy; FPLD, Dunnigan-type familial partial lipodystrophy; MAD, Mandibuloacral dysplasia; WRN, Werner syndrome. (C) The percentage of dysmorphic nuclei identified in U2OS cells transiently expressing the indicated Halo-lamin A mutants. Cells below a nuclear roundness cutoff of 0.82, as calculated in non-transfected cells, were marked as dysmorphic. Mean ± SEM; N = 3 experiments and > 50 cells per condition. ^**^ p < 0.01 by one-way ANOVA with Dunnett’s post-hoc test to wild type.

To characterize changes in dynamics of lamin A mutations by SMT, we transiently expressed the Halo-lamin A mutants in U2OS cells stably expressing LBR fragment-EGFP to mark the NE and analyzed molecules within the whole nucleus. While the majority of mutations associated with MAD and FPLD exhibited similar dynamics as wild type lamin A, several mutations associated with striated muscle myopathies (S22D, N195K, R50P) exhibited an increased diffusible population and faster dwell times (**Figure 4A, B**). Wild type lamin A had a diffusible population of 56% whereas S22D, N195K, R50P mutations increased this population to 68%, 68%, and 87% (p < 0.0001), respectively (**Figure 4A**). The fast and slow binding dwell times were reduced from 1.2 sec and 12 sec for wild-type lamin A to 1.1 sec and 9.4 sec for S22D, 1.1 sec and 9.8 sec for N195K, and 0.80 sec and 6.2 sec for R50P (p < 0.01 for all conditions) (**Figure 4B**). These results are in line with previously published studies in that the S22 residue has been shown to be phosphorylated during breakdown of the NE in mitosis [43] and during interphase to prevent proper lamin A incorporation into the lamina [44, 45]. The R50 residue mutation has been suggested to be critical for the proper polymerization of lamin A [46]. The depletion of the bound fraction of R50P to levels seen with Halotag alone serves as further validation that this fraction is reflective of lamin polymerization state. Intriguingly, mutations at S22 and N195 manifest as predominantly cardiac phenotypes and the R50P mutation has been linked to a severe form of EDMD with a cardiac phenotype [9, 47, 48].

**Figure 4:**
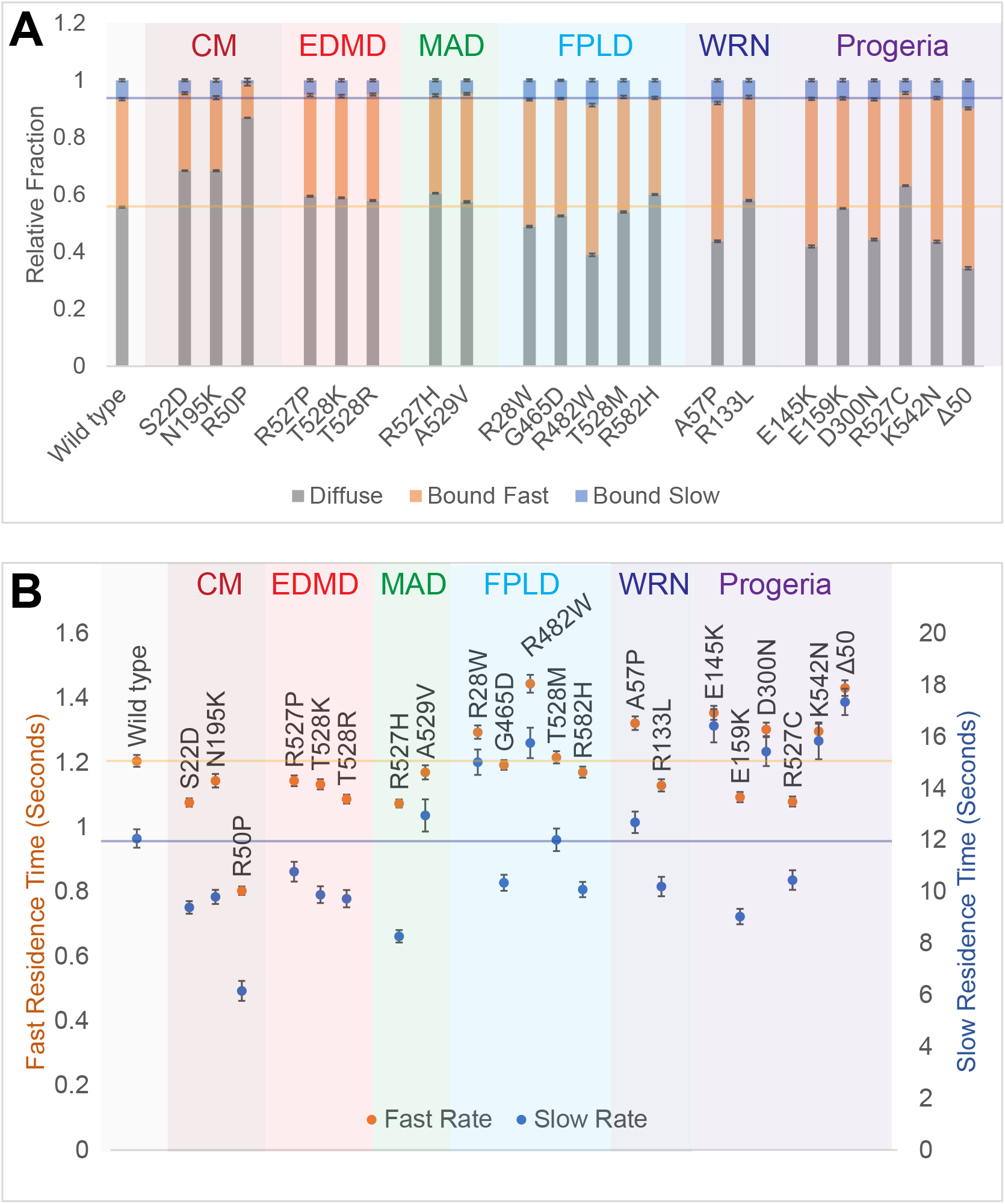
SMT of Halo-lamin A mutants. (A) Disease relevant mutations in lamin A were introduced and imaged by SMT as in Figure 1. Molecules within the nucleus, both at the periphery and in the nucleoplasm, were analyzed. Stacked bar graphs represent the ratio of diffuse (grey), fast bound (orange), and slow bound molecules (blue). Mutations are organized by respective disease outcome. Exposure time 30 ms; acquisition time 200 ms; 900 images per cell. CM, cardiomyopathy defect; EDMD, Emery-Dreifuss muscular dystrophy; FPLD, Dunnigan-type familial partial lipodystrophy; MAD, Mandibuloacral dysplasia; WRN, Werner syndrome. Values represent means and 95% confidence intervals of at least 10 cells and 2 experiments. (B) Fast bound (orange; left y-axis) and slow bound dwell times (blue; right y-axis) from molecules in (A).

Furthermore, mutations of R527 to P527 or T528 to K528 or R528 which are associated with slower progressing myopathies show statistically significant, but relatively minor, increases in the diffuse population of molecules over wild type (2–3%; p < 0.01), at most a 0.1 sec decrease in the fast bound dwell time (p < 0.01 for only the T528R mutant), and at most a 2.3 sec decrease in the slow bound dwell time (p < 0.01) (**Figure 4A, B**). Intriguingly, the overall smaller changes in binding properties in these mutations may provide insights as to why these mutations have thus far been predominantly linked to slowly progressing EDMD with minimal cardiac involvement.

In addition, the majority of the mutations associated with Werner syndrome and progeria (A57P, E145K, D300N, K542N, and Δ50) exhibited a consistent increase in the fast bound fraction by 10–14% for the point mutations and 18% for lamin Δ50 as well as longer fast and slow bound dwell times by 0.092–0.23 sec and 0.62–5.3 sec (p < 0.01 for all conditions) (**Figure 4A, B**). However, no statistically significant increase in the slow bound fraction was observed for the E145K, D300N nor K542N mutants (p > 0.05) (**Figure 4A**). Furthermore, an increase in the fast-bound fraction and dwell times was not consistent across all of the tested Werner- and progeria-linked mutations (R133L, E159K, R527C) (**Figure 4A, B**). These results suggest that some lamin A dynamics may be linked to disease type, especially with regards to cardiac dysfunction and systemic progeria. However, we do not find a consistent correlation between single molecule dynamics and specific types of laminopathies.

### 3.4. Analysis of distinct Halo-lamin A populations

Given the existence of a dynamic nucleoplasmic population of lamin A and a stable peripheral population, we reanalyzed our SMT data of the lamin mutants with separate ROIs to compare dynamic properties of molecules at the nuclear periphery (**Figure 5A**) to those of molecules in the interior of the nucleus (**Figure 5B**). Irrespective of the particular mutation, we find lamins at the nuclear periphery to have larger bound populations than lamin molecules in the nuclear interior (periphery: 29–65% bound vs. interior: 7–50%) (**Figure 5A**). At the periphery, the dwell times for the fast fraction were relatively consistent between mutations (1.0–1.4 sec), while the dwell times for the slow fraction exhibited the most variability (8.3–17 sec). In the nuclear interior, however, dwell times for both fast and slow fractions were highly variable between mutants (0.66–1.9 sec and 4.5–42 sec) (**Figure 5B**). The Halo-lamin Δ50 construct was not analyzed due to obfuscation from an increase in nuclear invaginations.

**Figure 5:**
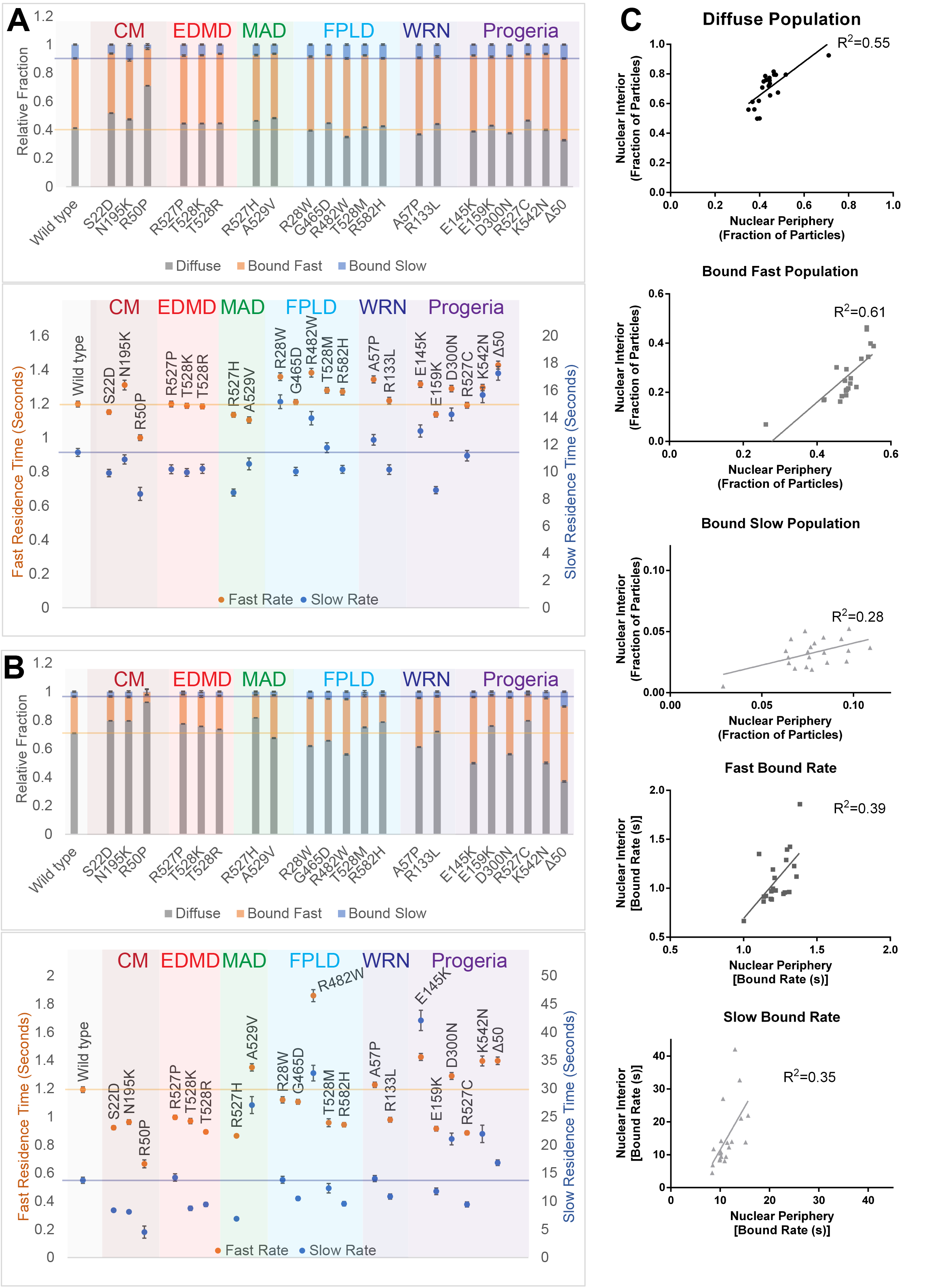
SMT of Halo-lamin A reveals compartment-specific differences in binding dynamics. (A) Relative ratios of diffuse, fast bound, and slow bound molecules (top) and the associated dwell times (bottom) of SMT data from Figure 4 where only the molecules located at the nuclear periphery were analyzed. (B) Relative ratios of diffuse, fast bound, and slow bound molecules (top) and associated dwell times (bottom) of SMT data from Figure 4 where only the molecules in the nuclear interior were analyzed. (C) The individual populations for each lamin mutation at the nuclear periphery from (A) plotted against the respective population in the nuclear interior from (B). Using each lamin mutant as a data point, linear regressions were fit to the relationship between nuclear compartments for each population: the ratio of diffuse, fast bound, and slow bound molecules as well as the fast and slow bound dwell times (top to bottom).

We next assessed whether lamin A mutations have disproportionate effects on molecules in the nuclear periphery versus the interior (**Figure 5C**). To do so, we plotted the dynamic properties of molecules at the nuclear periphery for each lamin mutation (from Figure 5A) against the respective properties of molecules in the nucleoplasm (from Figure 5B). If lamin mutations uniformly affected the dynamics of both populations, we would expect to see a well-fitting regression model. While the dwell times between the nuclear periphery and interior showed a high level of variance, the slow bound population showed the lowest correlation (**Figure 5C**), suggesting that mutations in lamin A differentially affect the stably bound population of lamin A at the nuclear periphery compared to the nucleoplasm, which consists of intranuclear lamin foci, whose function remains poorly understood [49]. Taken together, these results highlight the ability of SMT to distinguish and reveal population specific differences in lamin A behavior.

## 4. Discussion

Despite the prevalence of mutations in NE-associated proteins and their demonstrated relevance to human disease, it remains unknown how they cause the myriad of seemingly unrelated systemic and tissue-specific diseases referred to as laminopathies. Here, we developed SMT methodology for NE-associated proteins as a means to comprehensively characterize the dynamics of proteins at the nuclear periphery in a sensitive and accurate manner.

Lamin and NET dynamics have been indirectly assayed using solubility assays, nuclear elasticity, *in vitro* polymerization assays, and morphological changes, among others [32, 36, 42, 44-46, 50-54]. In contrast to these methods, SMT offers the advantages of direct quantification of protein behavior in the native cellular milieu [16]. SMT measurements reveal information about molecule localization, the types of binding behaviors, and the kinetics of these behaviors in a single experiment and at the single molecule level [16, 17]. Because each SMT experiment assays a large number of single molecules, individual molecules but also populations can be analyzed for complex behaviors unlike in bulk imaging methods such as FRAP and FCS. SMT is also uniquely suited to analyze NE-associating protein dynamics as it is able to distinguish the co-existing highly mobile and very stable populations of molecules. SMT can simultaneously extrapolate information about both short- and long-term binding events, which is technically challenging and requires imaging over long intervals when using FRAP or FCS. However, SMT is limited in that it typically relies on exogenous protein expression. One approach to circumvent this limitation is the use of CRISPR methodology to tag endogenous proteins [55]. In addition, while molecule tracking offers snapshots of protein behavior in native environments, the acquired data can vary with imaging setup and acquisition time, making it difficult to translate data across studies [18].

We show here that SMT is able to accurately measure and distinguish dynamics of different lamin variants as well as the transmembrane protein, Samp1. As evidence for the specificity of the method, the behavior of lamins and NETs is distinct from Halotag alone and treatment of lamin A Δ50 with an FTI was able to significantly reduce the protein’s bound fraction and dwell times. Furthermore, SMT of a panel of lamin A mutants revealed the technique to be sensitive enough to profile mutation-specific effects on binding dynamics that were not observed by morphology alone. Of note, we identified three mutations (S22D, N195K, R50P) with altered binding dynamics. Interestingly, these loss-of-function mutations are linked to early onset and severe striated muscle dysfunction with a cardiac phenotype, which is reminiscent of the effect of *LMNA* knockout in mice [56]. In contrast, we observed increased binding in a number of progeria-linked mutations (A57P, E145K, D300N, K542N, and Δ50), again, in line with the observed increased retention of lamin Δ50 [25, 42], possibly pointing to a link between reduced dynamics and systemic pre-mature aging.

We did not observe disease-specific patterns of behavior with the other EDMD-, MAD-, and FPLD-linked mutations, suggesting that these mutations may have a separate etiology and/or there are cell-type specific nuances that need to be considered. In the case of slowly progressing EDMD, it is possible that the changes in dynamics are small and require years of strain to progress to a clinical manifestation. Another hypothesis is that these mutations differentially alter the dynamics of other NE-associated proteins, such as emerin [52]. Furthermore, we did not observe notable changes in dynamics between different mutations occurring at the same residue (R527 and T528) despite their links to different diseases nor did we note consistent effects in mutations in the same protein domain. Hence, the changes in lamin dynamics may be largely dependent on the specific residue that is altered, and we conclude that changes in dynamics are not the only determinant of disease. Indeed, the single residue mutations in *LMNA* have been shown to differentially alter protein-protein interactions [57]. Nevertheless, these proof-of-concept experiments highlight that SMT of nuclear lamins and NETs as a robust, sensitive, and easily adaptable system to profile protein dynamics at the nuclear periphery.

Taken together, our data establish SMT as a sensitive means to profile lamin and NETs dynamics. We suggest that SMT will be a useful tool to directly test how extrinsic factors such as pharmacological interventions, extracellular matrix stiffness, cytoskeleton stability, binding partner knockdown, as well as intrinsic protein properties, including mutations or polymerization state, influence the dynamic behavior of disease-related mutants of lamins and NETs.

## Acknowledgments

This research was supported by the Intramural Research Program of the National Institutes of Health, National Cancer Institute, and Center for Cancer Research. High-throughput imaging work was performed at the High-Throughput Imaging Facility (HiTIF)/Center for Cancer Research/National Cancer Institute/NIH.

---

**Supplementary Table 1:**
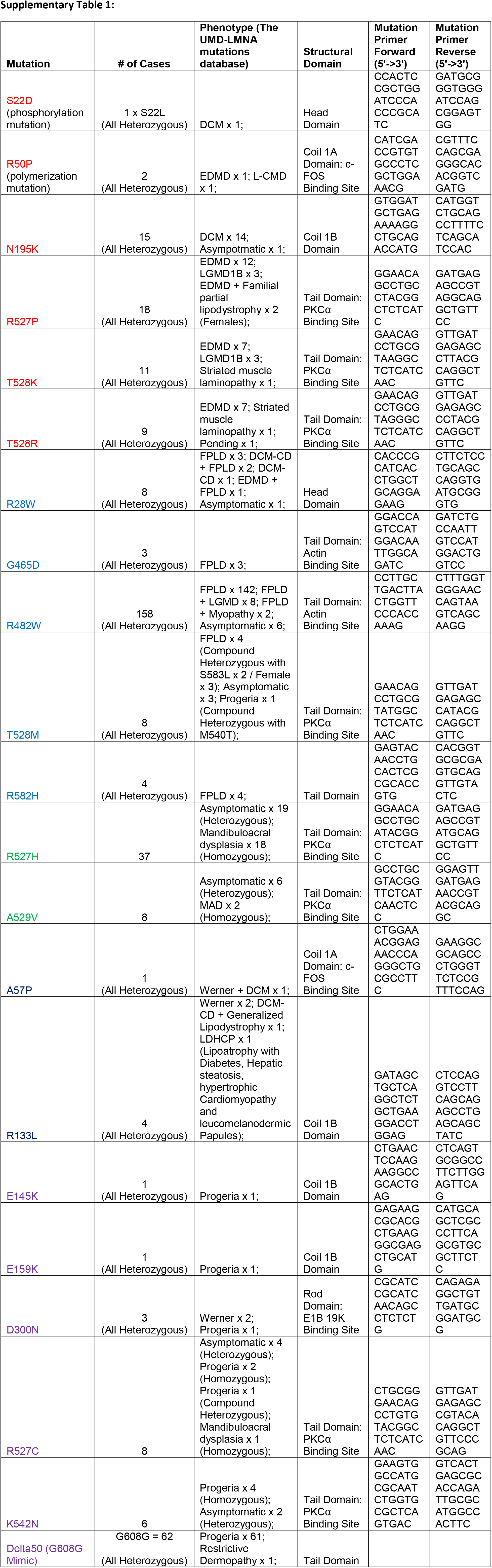

